# VISTA as a ligand downregulates LPS-mediated inflammation in macrophages and neutrophils

**DOI:** 10.1101/2021.10.17.464340

**Authors:** Yu-Heng Vivian Ma, Amanda Sparkes, Jean Gariépy

## Abstract

V-domain immunoglobulin suppressor of T-cell activation (VISTA) has emerged as a unique immunoregulatory receptor on cells of the myeloid lineage. Agonizing VISTA on myeloid cells has recently been demonstrated to have a profound effect on dampening inflammatory responses. VISTA has been proposed to function both as a ligand and as a receptor. In this context, the role of VISTA as a ligand has been largely ignored. Using a high-avidity agonist of the VISTA receptor (VISTA-COMP), we investigated the effect of exogenous VISTA, as a ligand, on macrophages and neutrophil cellular pathways in an acute inflammatory setting. RNA sequencing analysis demonstrated that VISTA-COMP downregulates pro-inflammatory cytokines and chemokines and upregulates immunoregulatory genes in both LPS-stimulated macrophages and neutrophils ex vivo. Interestingly, unlike VISTA itself, the receptor is only expressed following LPS stimulation of these cell populations. Furthermore, the administration of VISTA-COMP attenuated the rise in circulating TNFα levels in LPS-treated mice in vivo. These results suggest that VISTA serves a redundant role on macrophages and neutrophils acting as both a ligand and a receptor in the context of an acute inflammatory event.

## Introduction

V-domain immunoglobulin suppressor of T-cell activation (VISTA), a member of the immunoglobulin superfamily, is described as an immune checkpoint molecule regulating T-cell activation. It is widely expressed within the hematopoietic compartment with the highest expression being found on myeloid cells, particularly macrophages (1–5). Mechanistically, VISTA has been proposed to function both as a ligand and as a receptor (1,2,4–8) with the majority of published studies focusing on the differential expression of VISTA on myeloid cells and the consequences of agonizing or antagonizing VISTA itself. In particular, Bharai and co-workers reported that the overexpression of VISTA on monocytes enhanced cytokine secretion (9); while blocking VISTA on MDSCs mitigates their suppressive activity (4). More recently, VISTA has been found to have a profound impact on chemotaxis (10); and agonizing VISTA on myeloid cells dampens inflammatory immune responses (7) (reviewed by ElTanbouly and Zhao *et al*. (11)). None of the aforementioned studies examine the role of a VISTA receptor on myeloid cells. This is due, in part, on the lack of a definitive receptor identified for VISTA on murine myeloid cells. To facilitate VISTA receptor activation studies in the absence of Fc-related events, we previously developed a multimeric recombinant form of the extracellular domain of murine VISTA, termed VISTA-COMP (12). This construct was engineered by fusing the IgV domain of VISTA to a short peptide corresponding to the pentamerization domain of the cartilage oligomeric matrix protein. The resulting VISTA pentamer proved superior to VISTA-Fc in terms of binding to the VISTA receptor on murine T-cells and as a soluble factor able to dampen their proliferation and the production of pro-inflammatory cytokines (12). Here, we investigated the mechanistic impact of exogenously treating myeloid immune cell populations with VISTA-COMP in an acute inflammatory setting, namely upon stimulating them *ex vivo* with lipopolysaccharides (LPS) in the context of a murine endotoxemia model. As outlined in the present study, RNA sequencing analyses demonstrated that VISTA-COMP downregulates the expression of pro-inflammatory cytokines and chemokine and upregulates immunoregulatory genes in both LPS-stimulated macrophages and neutrophils *ex vivo*. Furthermore, we showed that the administration of VISTA-COMP *in vivo* attenuates the characteristic monophasic spike in circulating TNFα levels (13,14) in mice treated with LPS. Overall, both agonistic antibody binding to VISTA (7) and VISTA itself binding to its receptor on activated macrophages or neutrophils (this study) trigger similar cell signaling pathways leading to a rapid dampening of myeloid cell-triggered pro-inflammatory responses.

## Methods

### Mice

Female C57BL/6 mice at 8–10 weeks of age (The Jackson Laboratory, Bar Harbor, ME, USA) were used throughout this study and housed at the Comparative Research Facility at Sunnybrook Research Institute (SRI; Sunnybrook Health Sciences Centre, Toronto, ON, Canada). All protocols were approved by the SRI Comparative Research Animal Care Committee, accredited by the Canadian Council of Animal Care.

### Macrophage and neutrophil isolation

Brewer thioglycolate medium (3%, 4 mL, autoclaved; MilliporeSigma, St. Louis, MO, USA) was injected intraperitoneally into mice to recruit macrophages and neutrophils to the peritoneal cavity. Mice were sacrificed either 1 day or 4 days later and peritoneal exudates cells (PECs) were harvested by peritoneal lavage with PBS. PECs recovered after 4 days consisted mostly of macrophages, whereas PECs isolated 1 day after thioglycolate injection consisted mostly of neutrophils (15). The identity of the PECs was confirmed using flow cytometry by staining them with the following antibodies: APC/Cy7 anti-CD45, BV510 anti-CD11b, PE/Cy7 anti-Ly6G, Alexa Fluor 700 anti-Siglec F, mCherry anti-Ly6C, Pacific Bule anti-IA/IE, PerCP/Cy5.5 anti-CD11c, and FITC anti-F4/80 (BioLegend, San Diego, CA, USA). Binding of recombinant mouse VISTA-COMP (12) was validated by incubating the isolated PECs with biotinylated VISTA-COMP or COMP (produced in house) at 4 °C for 1 hour, followed by detection with a PE streptavidin conjugate (BioLegend). Stained PECs were detected using a BD LSR II flow cytometer (BD, Franklin Lakes, NJ, USA) maintained by The Centre for Flow Cytometry & Scanning Microscopy (CCSM) at SRI. Throughout the study, PECs were maintained at 37 °C and 5% CO_2_ in RPMI 1640 medium (Wisent, St-Bruno, QC, Canada) supplemented with 10% heat-inactivated fetal bovine serum (FBS) (Wisent), 1% penicillin-streptomycin (P/S) (Wisent), 0.05mM 2-mercaptoethanol (MilliporeSigma), and 20mM HEPES (Wisent).

### RNA sequencing

Neutrophils and macrophages (PECs isolated one day or 4 days post-thioglycolate injection, respectively) were stimulated with 1 µg/mL lipopolysaccharides (LPS; *Escherichia coli* O111:B4; MilliporeSigma) and 15 µg/mL VISTA-COMP for 2 hours. RNA content was isolated from the conditioned macrophages or neutrophils using the RNeasy Mini Kit (Qiagen, Venlo, Netherlands), treated with DNA-free Kit (Life Technologies, Carlsbad, CA, USA), and sent in for RNA-sequencing at The Donnelly Sequencing Centre (University of Toronto). The library was prepared with TruSeq Stranded mRNA Library Prep Kit (Illumina, San Diego, CA, USA) and sequenced with NextSeq500 (High Output, 75 Cycles, v2 Chemistry; R1: 85bp, IR1: 6bp, single read) (Illumina). This sequencing data have been deposited in the ArrayExpress database at EMBL-EBI (www.ebi.ac.uk/arrayexpress) under accession number E-MTAB-10856. Read counts were obtained with RNA Express on Illumina’s BaseSpace (https://basespace.illlumina.com), which aligns RNA-sequencing reads to reference mouse genome mm10 (GRCm38) (16) using STAR (17). Differential gene expression was computed using DESeq2 v1.26.0 in R/Bioconductor (18). Enrichment of gene ontology terms (19,20) were computed on the significantly upregulated and downregulated genes using GOseq v1.38.0 (21) in R/Bioconductor with count bias accounted for. Revigo (22) was used to trim own the list of gene ontology terms. Heatmaps were prepared with edgeR v3.28.1 (23) and gplots v3.0.4 (24) in R/Bioconductor.

### qPCR and cytokine analysis

Neutrophils and macrophages (PECs isolated one day or 4 days post-thioglycolate injection, respectively) were stimulated with 1 µg/mL LPS and 15 µg/mL VISTA-COMP. For qPCR, RNA contents were isolated from the cells 1 or 2 hours after stimulation using TRIzol Reagent (Thermo Fisher Scientific) and reverse-transcribed using high-capacity cDNA reverse transcription kit (Thermo Fisher Scientific). qPCR was performed with the SensiFAST SYBR no-ROX kit (Meridian Bioscience, Cincinnati, OH, USA) using gene-specific primers listed in Table 1 (Integrated DNA Technologies, Coralville, IA, USA). For cytokine analysis, supernatants were harvested 3 hours after stimulation and the concentrations of TNFα, IL-6, and CXCL1 in each supernatant were assessed by flow cytometry (BD LSR II; Sunnybrook Health Sciences Centre) using the LegendPlex kit (BioLegend).

**Table 1:**
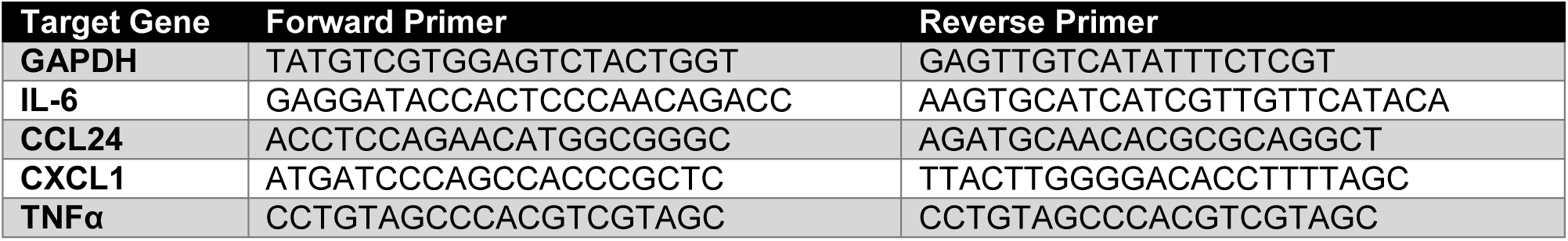
Gene-specific primers used in qPCR.

### Endotoxemia

LPS (10 mg/kg) and VISTA-COMP (30 mg/kg) were injected intraperitoneally into mice. After 90 minutes, mice were sacrificed, and blood was collected by cardiac puncture. Serum concentrations of TNFα were measured by flow cytometry (BD LSR II flow cytometer) using the LegendPlex kit (BioLegend).

### ELISA

To confirm non-homotypic binding of VISTA, an ELISA was performed. ELISA plate wells (Thermo Fisher Scientific) were coated with VISTA-Fc (1 µg/ml; produced in house) and subsequently blocked with 0.5% milk. VISTA-COMP (5 µg/ml) was then dispensed into wells, and incubated for 1 hour at room temperature. Binding was detected using an anti-his HRP conjugated antibody (MilliporeSigma), and the substrate TMB (3,3’,5,5’-tetramethylbenzidine; Thermo Fisher Scientific). As a positive control for the ELISA, PDL1-COMP binding PD1-Fc was also performed as described. Furthermore, due to homology, VISTA-COMP binding activity against PD1-Fc was also assessed. Absorbance readings were recorded at 450 nm.

### Statistics

Each experiment was performed with a minimum of 3 biological replicates, where each replicate represented one mouse. Experiments using PECs were performed by splitting PECs isolated from one mouse into different stimulation groups to allow for paired analysis. Student’s t-test (2-tail, paired for experiments involving PECs) was used to test significance between different stimulation groups (significance was taken at α = 0.05). For RNA sequencing results, a gene is considered to be differentially expressed when the adjusted p-value (corrected for multiple testing with the Benjamini-Hochberg method) calculated by DESeq2 is less than 0.05.

## Results

### VISTA downregulates the inflammatory immune response and chemotaxis in LPS-stimulated macrophages

VISTA had been proposed to act as both a ligand and a receptor. It has been recently demonstrated that during LPS-induced inflammation, VISTA acts as a receptor to reprogram macrophages (7). In the present study, we investigated instead the role of VISTA as a ligand binding to a VISTA receptor on LPS-activated macrophages. A high avidity form of the extracellular IgV domain of VISTA (VISTA-COMP) served as our ligand as it behaves as a true VISTA receptor agonist that lacks any Fc-related functions associated with the use of antibodies and VISTA-Fc constructs (12).

Macrophages were isolated as peritoneal exudates cells (PECs) from mice 4 days after thioglycolate injection (15). Their macrophage identity was confirmed by flow cytometry (Supplementary Figure 1). The isolated macrophages were stimulated with PBS, LPS alone (1 µg/mL LPS), LPS/VISTA (1 µg/mL LPS and 15 µg/mL VISTA-COMP), or VISTA alone (15 µg/mL VISTA-COMP) for 2 hours, followed by RNA sequencing. Sets of genes that displayed the largest differences in expression between macrophages stimulated with LPS and those stimulated with LPS/VISTA were then classified into distinct biological processes. Specifically, heat maps were constructed where significantly differentially expressed genes (red: high expression; blue: low expression) were grouped under different biological processes, and ranked based on the difference in gene expression between macrophages stimulated with LPS and LPS/VISTA (**Figure 1**). Notably, genes that were found to be significantly differentially expressed when comparing LPS- to LPS/VISTA-stimulated macrophages were not found to be significantly different in the absence of LPS stimulation (PBS-VS VISTA-stimulated). This finding suggests the VISTA receptor is only present on macrophages following the LPS stimulation, in contrast to VISTA which is expressed on non-stimulated Thio-PECS (25). This conclusion was confirmed by flow cytometry where VISTA-COMP binding to macrophages was only observed following LPS stimulation (Supplementary Figure 1). As shown in **Figure 1**, biological processes involved in immune responses, such as inflammation and chemotaxis, are downregulated in macrophages treated with LPS/VISTA as compared to the LPS only group. Importantly, *il6*, a hallmark cytokine produced in response to LPS stimulation (26), is shown to be significantly reduced in the LPS/VISTA-treated group. Furthermore, *ccl24* and *cxcl1*, genes involved in multiple immune- and chemotaxis-related biological processes, were downregulated by VISTA in the LPS-stimulated macrophages. The levels of expression of these specific genes were subsequently verified by qPCR on additional biological replicates and shown to be significantly lower in the LPS/VISTA-as compared to the LPS-treated group (*il6* by 30%; *ccl24* by 27%; *cxcl1* by 10%) (**Figure 2**). Moreover, although a downregulation in *tnf* was not observed in LPS-stimulated macrophages after 2 hours of stimulation with VISTA-COMP, it has been shown by others to typically precedes *il6* production in a standard response to LPS (13,14). Therefore, we re-assessed *tnf* expression in macrophages at an earlier time-point, namely, after 1 hour of stimulation. As expected, we found that *tnf* expression was significantly reduced by 56% in macrophages stimulated by LPS/VISTA as compared to those stimulated by LPS alone (**Figure 2**). To confirm that these changes in gene expression did result in changes in cytokine secretion, we measured the concentration of selected cytokines in the supernatant of stimulated macrophages, and found that concentrations of TNFα, IL-6, and CXCL1 were lower in macrophages exposed to LPS/VISTA in comparison to macrophages that were exposed to LPS only (TNFα by 28%; IL-6 by 36%; CXCL1 by 30%) (**Figure 3**).

**Figure 1:**
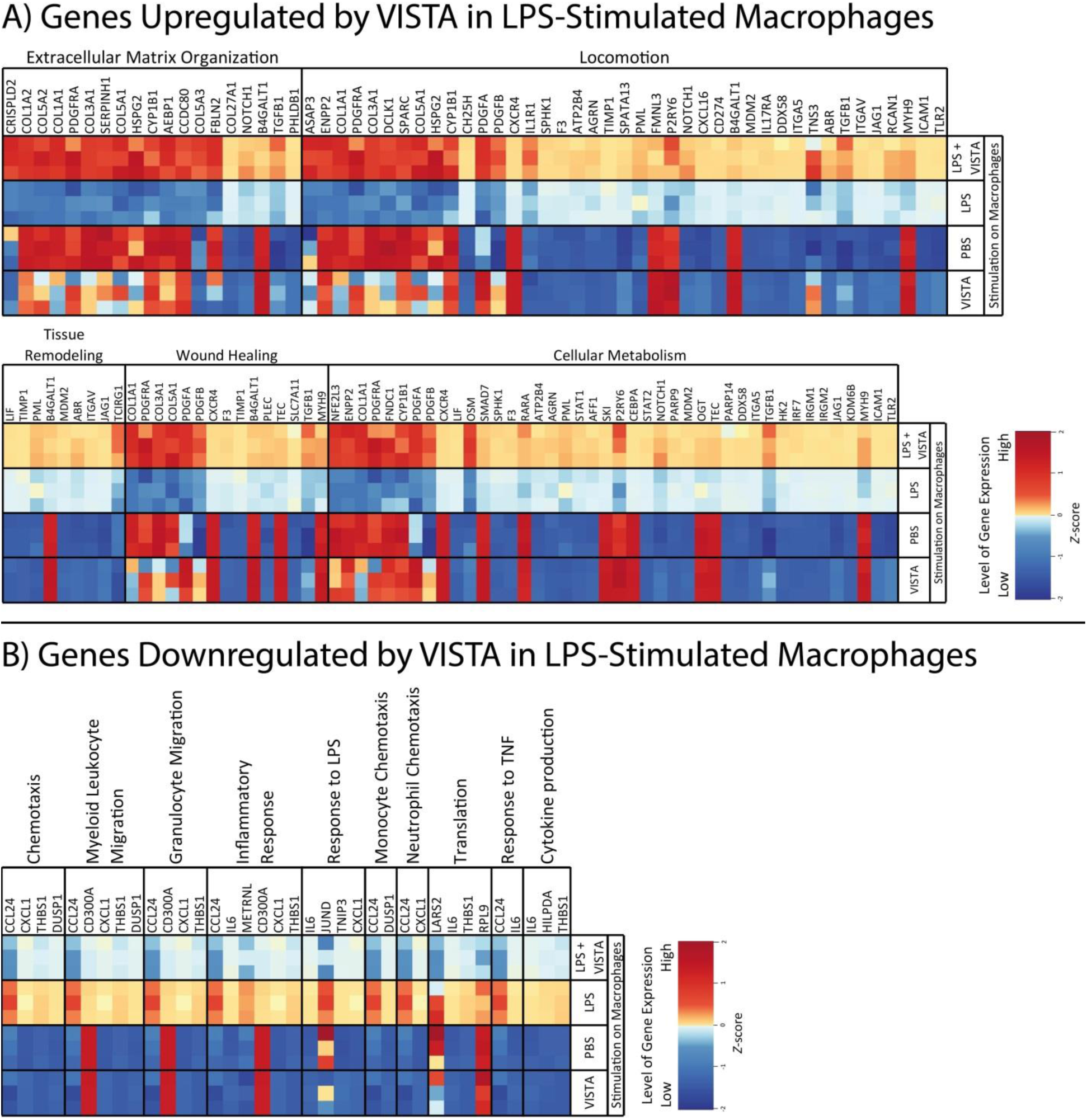
Differential gene expression between macrophages stimulated *ex vivo* with LPS only or LPS and VISTA. RNA sequencing was performed on macrophages (PECs isolated 4 days after thioglycolate injection) stimulated with PBS, LPS (1 µg/mL LPS), LPS and VISTA (15 µg/mL VISTA-COMP), or VISTA for 2 hours (n = 3 shown as individual squares on the heat map). Heat map (red: high expression; blue: low expression) was shown on selected genes that were significantly (adjusted *p*-value < 0.05) **(A)** upregulated or **(B)** downregulated by VISTA in LPS-stimulated macrophages, grouped by biological processes these genes are involved in.

**Figure 2:**
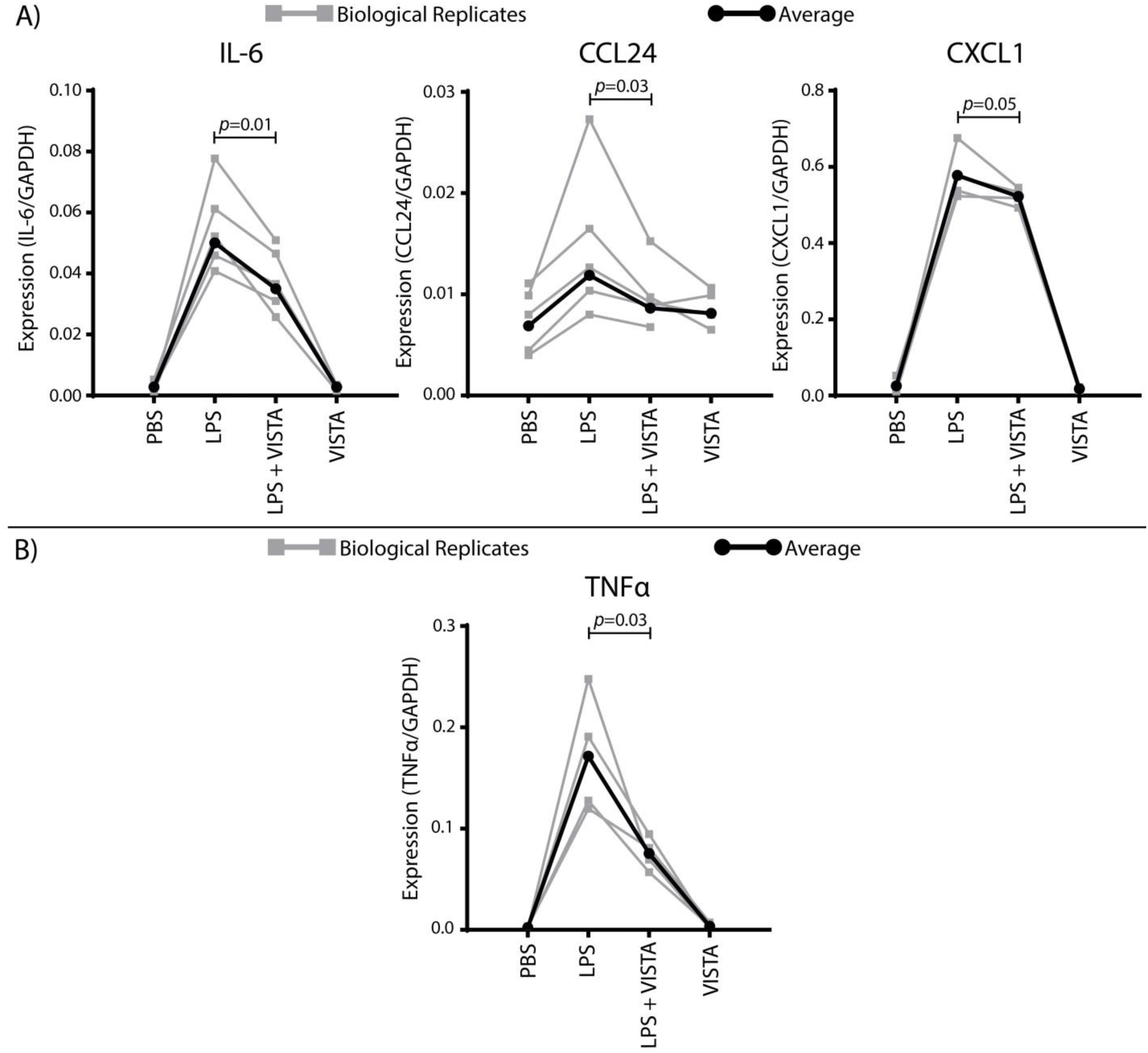
VISTA downregulates macrophage production of key genes stimulated by LPS. qPCR showing the gene expression (normalized to GAPDH expression) by macrophages (PECs isolated 4 days after thioglycolate injection) stimulated with PBS, LPS (1 µg/mL LPS), LPS and VISTA (15 µg/mL VISTA-COMP), or VISTA for **(A)** 2 hours or **(B)** 1 hour. Each grey line represents an experiment performed on macrophages isolated from a single mouse. Black lines represent the average of all experiments.

**Figure 3:**
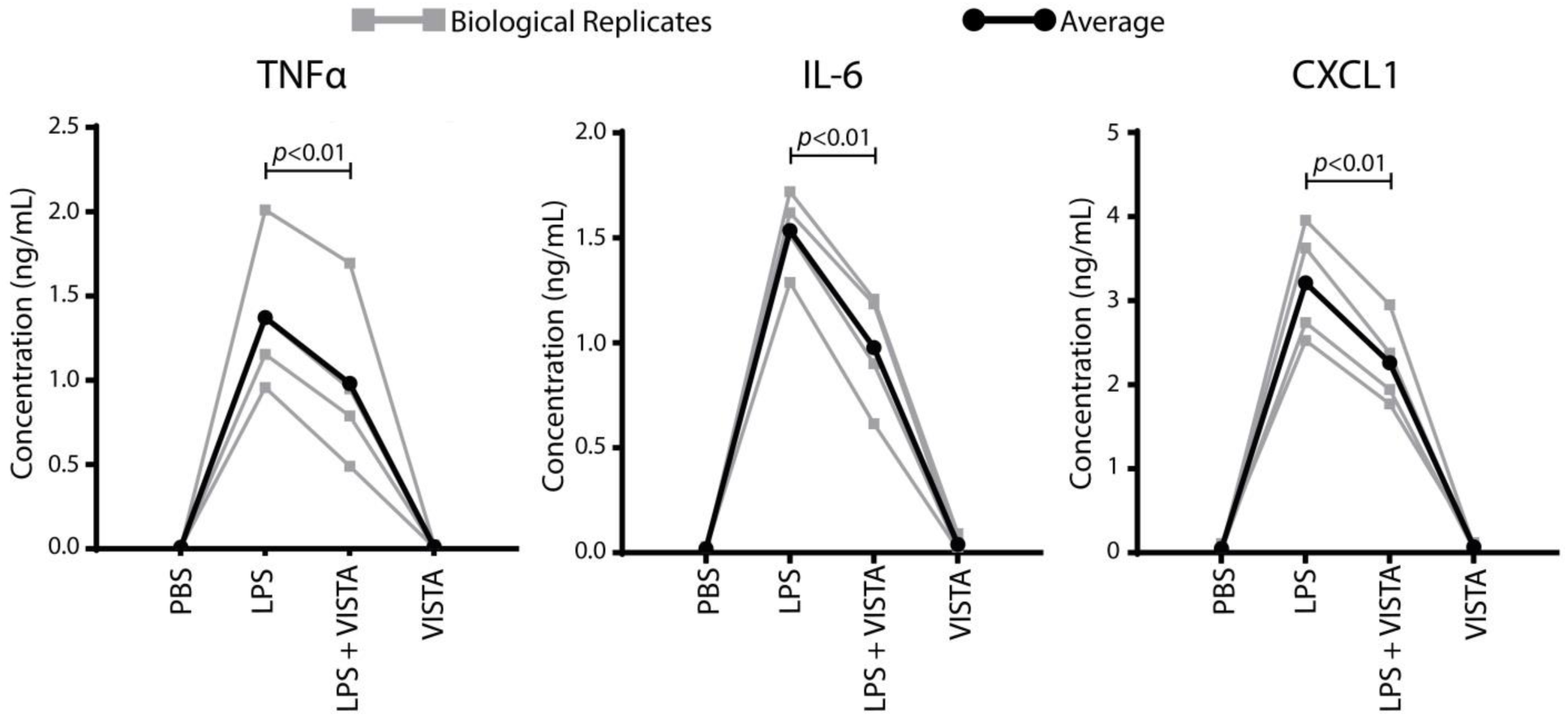
VISTA downregulates secretion of key cytokines by macrophage stimulated by LPS. Concentration of cytokines in the supernatant of macrophages (PECs isolated 4 days after thioglycolate injection) stimulated with PBS, LPS (1 µg/mL LPS), LPS and VISTA (15 µg/mL VISTA-COMP), or VISTA for 3 hours. Each grey line represents an experiment performed on macrophages isolated from a single mouse. Black lines represent the average of all experiments.

A greater number of genes were upregulated in LPS-stimulated macrophages by VISTA-COMP relative to the macrophages treated with LPS only (**Figure 1**). Importantly, the most represented biological processes associated with this set of upregulated genes were not linked to inflammatory or immune responses. Notably, genes associated with tissue remodeling and wound healing pathways were upregulated by VISTA in the LPS-stimulated macrophages, suggesting a reprogramming of these macrophages to a M2 subtype (27). This conclusion is further supported by the upregulation of *tgfb1* (27). The upregulation of LIF also suggests an immunoregulatory subtype of macrophages (28). Interestingly, although chemotactic genes were downregulated by VISTA in LPS-stimulated macrophages, genes involved in general locomotion appear to be upregulated. This finding may reflect an upregulation of biological processes such as extracellular matrix organization, wound healing, and tissue remodeling, that require locomotion. In particular, *cxcr4*, a receptor involved in cell migration and known for its role in reprogramming macrophages into a M2 subtype in tumor microenvironment (29), was found to be upregulated in the LPS/VISTA-stimulated macrophages, as compared to the LPS-stimulated macrophages. Finally, several genes involved in metabolism were also found to be upregulated by VISTA in the LPS-stimulated macrophages. Overall, VISTA-COMP (this study) and anti-VISTA antibody (7) affect the gene expression of LPS-stimulated macrophages in remarkably similar ways. Binding ELISA between VISTA-COMP and VISTA-Fc shows that VISTA does not bind in a homotypic fashion (Supplementary Figure 2), suggesting that signaling through VISTA and the VISTA receptor are two redundant but distinct pathways.

### VISTA downregulates the inflammatory immune response and chemotaxis in LPS-stimulated neutrophils

In addition to macrophages, neutrophils represent another important cell population involved in LPS response (30). Furthermore, to our knowledge, there is currently no data available with regard to the influence of exogenous VISTA signaling through a receptor on neutrophils. Therefore, we proceeded to perform the same gene ontology studies as described above on neutrophils. PECs isolated from mice 1 day after thioglycolate injection were used as neutrophils (15), and their identity was confirmed using flow cytometry (Supplementary Figure 3). The neutrophils were stimulated with nothing (PBS), LPS (1 µg/mL LPS), LPS/VISTA (1 µg/mL LPS and 15 µg/mL VISTA-COMP), or VISTA (15 µg/mL VISTA-COMP) for 2 hours and gene expression levels assessed by RNA sequencing. Heat maps were constructed to highlight genes exhibiting significant differences in expression between LPS- and LPS/VISTA-stimulated neutrophils (red: high expression; blue: low expression; ranked by magnitude of difference between LPS- and LPS/VISTA-stimulated neutrophils). These genes were then regrouped under specific biological processes. As was the case with macrophages, VISTA only affected the expression of genes in LPS-treated neutrophils, and VISTA-COMP only bound to neutrophils that were stimulated with LPS (Supplementary Figure 3).

To investigate whether neutrophils responded to VISTA in a similar manner to macrophages following LPS stimulation, we first confirmed that key genes found to be downregulated by VISTA in LPS-stimulated macrophages were also downregulated by VISTA in LPS-stimulated neutrophils. As projected, the expression of both *il6* and *cxcl1* genes were found to be significantly lower in LPS/VISTA-stimulated neutrophils as compared to LPS-stimulated neutrophils (**Figure 4**) and confirmed by qPCR (*il6* by 27 %; *cxcl1* by 38%) (**Figure 5**). The level of gene expression of *tnf* was also downregulated by VISTA in LPS-stimulated neutrophils at this 2-hour time point (**Figure 4**). However, to be consistent with the time dependency of gene expression profiles observed in macrophages, neutrophils were also stimulated with PBS, LPS, LPS/VISTA, and VISTA for 1 hour, and measured the level of *tnf* gene expression. At this time point, the expression of *tnf* was significantly reduced (by 67%) in neutrophils stimulated with LPS/VSITA relative to neutrophils stimulated with LPS alone (**Figure 5**). In addition, several other genes involved in immune-related biological processes, such as the differentiation and chemotaxis of myeloid cells and lymphocytes in general, were found to be significantly downregulated by VISTA in LPS-stimulated neutrophils as well. All these results point to a pathway where VISTA signals neutrophils to downregulate their inflammatory response to LPS.

**Figure 4:**
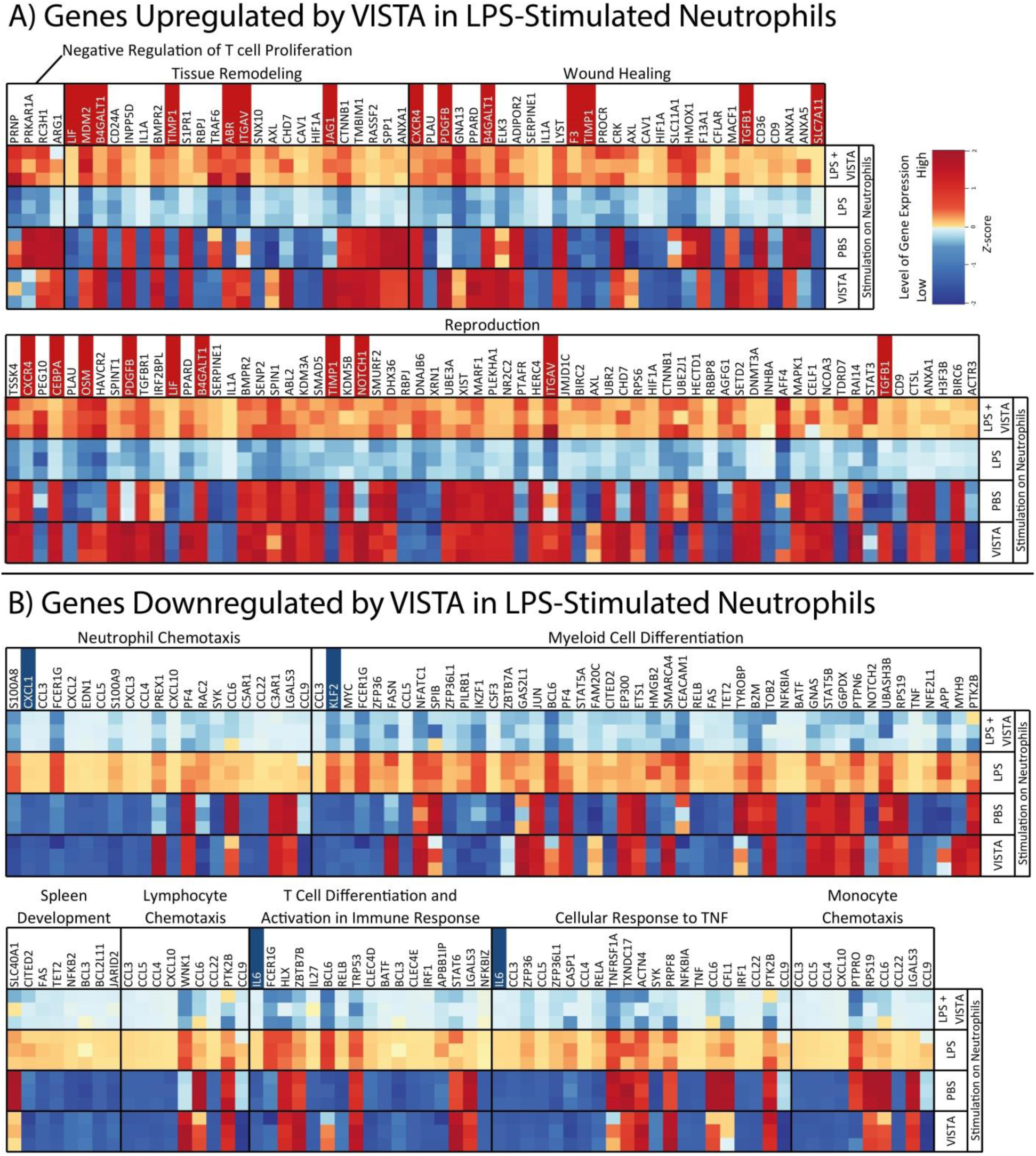
Differential gene expression between neutrophils stimulated *ex vivo* with LPS only or LPS and VISTA. RNA sequencing was performed on neutrophils (PECs isolated 1 day after thioglycolate injection) stimulated with PBS, LPS (1 µg/mL LPS), LPS and VISTA (15 µg/mL VISTA-COMP), or VISTA for 2 hours (n = 3 shown as individual squares on the heat map). Heat map (red: high expression; blue: low expression) was shown on selected genes that were significantly (adjusted *p*-value < 0.05) **(A)** upregulated or **(B)** downregulated by VISTA in LPS-stimulated neutrophils, grouped by biological processes these genes are involved in. Gene names highlighted in red are genes significantly upregulated by VISTA in both LPS-stimulated macrophages and neutrophils; gene names highlighted in blue are genes that were significantly downregulated by VISTA in both LPS-stimulated macrophages and neutrophils (Figure 2).

**Figure 5:**
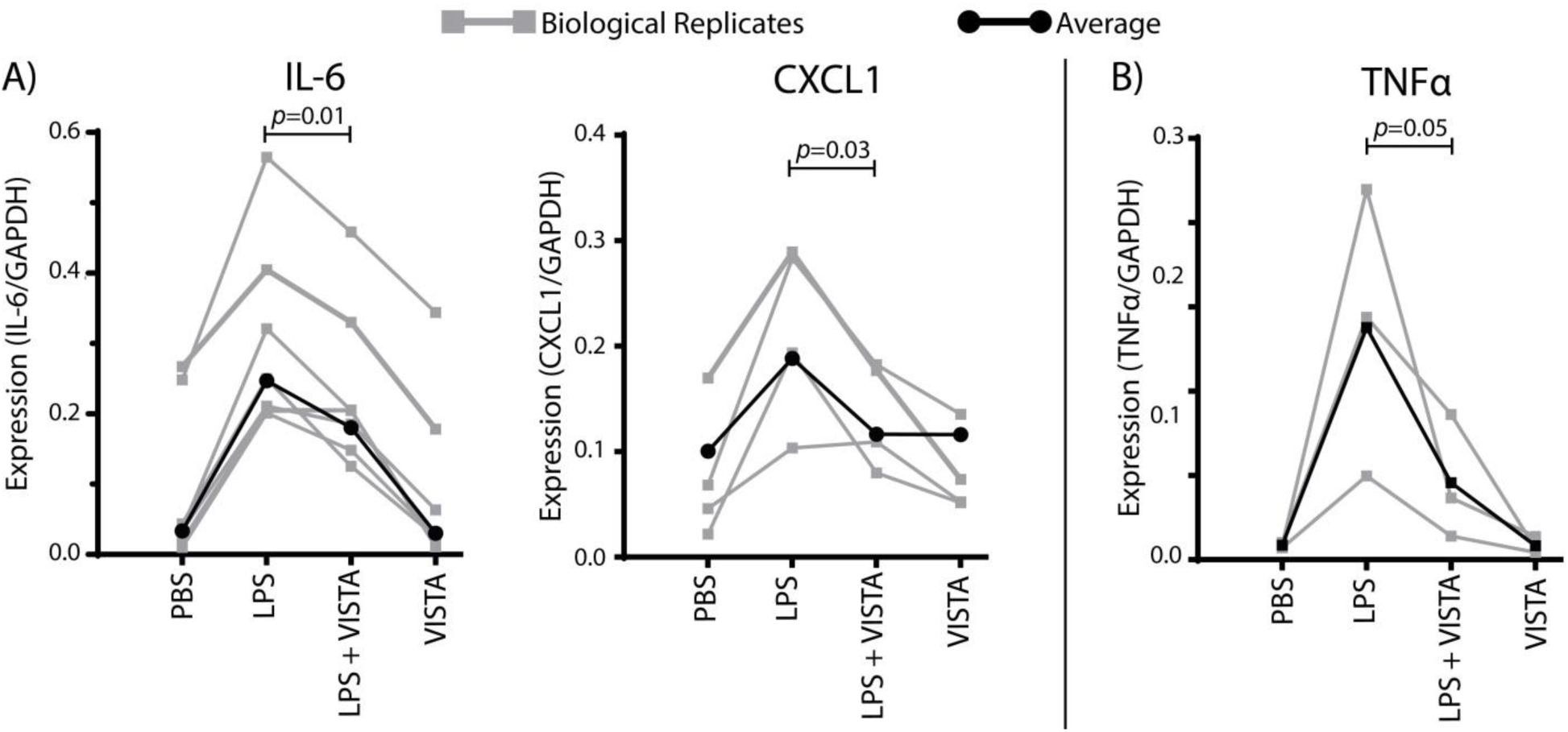
VISTA downregulates neutrophil production of key genes stimulated by LPS. qPCR showing the gene expression (normalized to GAPDH expression) by neutrophils (PECs isolated 1 day after thioglycolate injection) stimulated with PBS, LPS (1 µg/mL LPS), LPS and VISTA (15 µg/mL VISTA-COMP), or VISTA for **(A)** 2 hours or **(B)** 1 hour. Each grey line performed on neutrophils isolated from a single mouse. Black lines represent the average of all experiments.

The biological processes that were upregulated by VISTA in LPS-stimulated neutrophils were similar to those observed in LPS/VISTA-stimulated macrophages. In particular, genes linked to wound healing and tissue remodeling, along with key mediators of anti-inflammatory subtypes *lif* (28), *tgfb1* (27), and *cxcr4* (29), were also upregulated by VISTA in LPS-stimulated neutrophils. Interestingly, a set of genes upregulated in the LPS/VISTA group compared to the LPS group were found to be involved in reproduction. This classification may reflect genes that are also involved in other upregulated functions, such as locomotion and metabolism. In addition, many genes involved in pregnancy are also associated with anti-inflammatory and immune modulatory functions, such as *lif, tgfb1*, and *osm* (31). Overall, these results suggest that VISTA acts similarly on LPS-stimulated neutrophils to promote an immune-regulatory response.

### VISTA reduces the monophasic serum TNFα peek in a mouse model of endotoxemia

Given the suppressive activity of VISTA-COMP *ex vivo* in an acute inflammatory setting of LPS stimulation, we assessed its efficacy in the context of an *in vivo* model. Endotoxemia is a well-established *in vivo* model to assess therapeutic response following LPS insult. Since increase in TNFα is a hallmark of sepsis (26), treatments leading to a reduction of serum TNFα levels has been shown to protect mice against LPS-induced lethality (32). As such, we measure the circulating levels of TNFα in the serum of mice treated with PBS, LPS, or LPS and VISTA-COMP. As shown in **Figure 6**, serum levels of TNFα were reduced by 49% in animals treated with VISTA-COMP and LPS relative to LPS only-treated mice.

**Figure 6:**
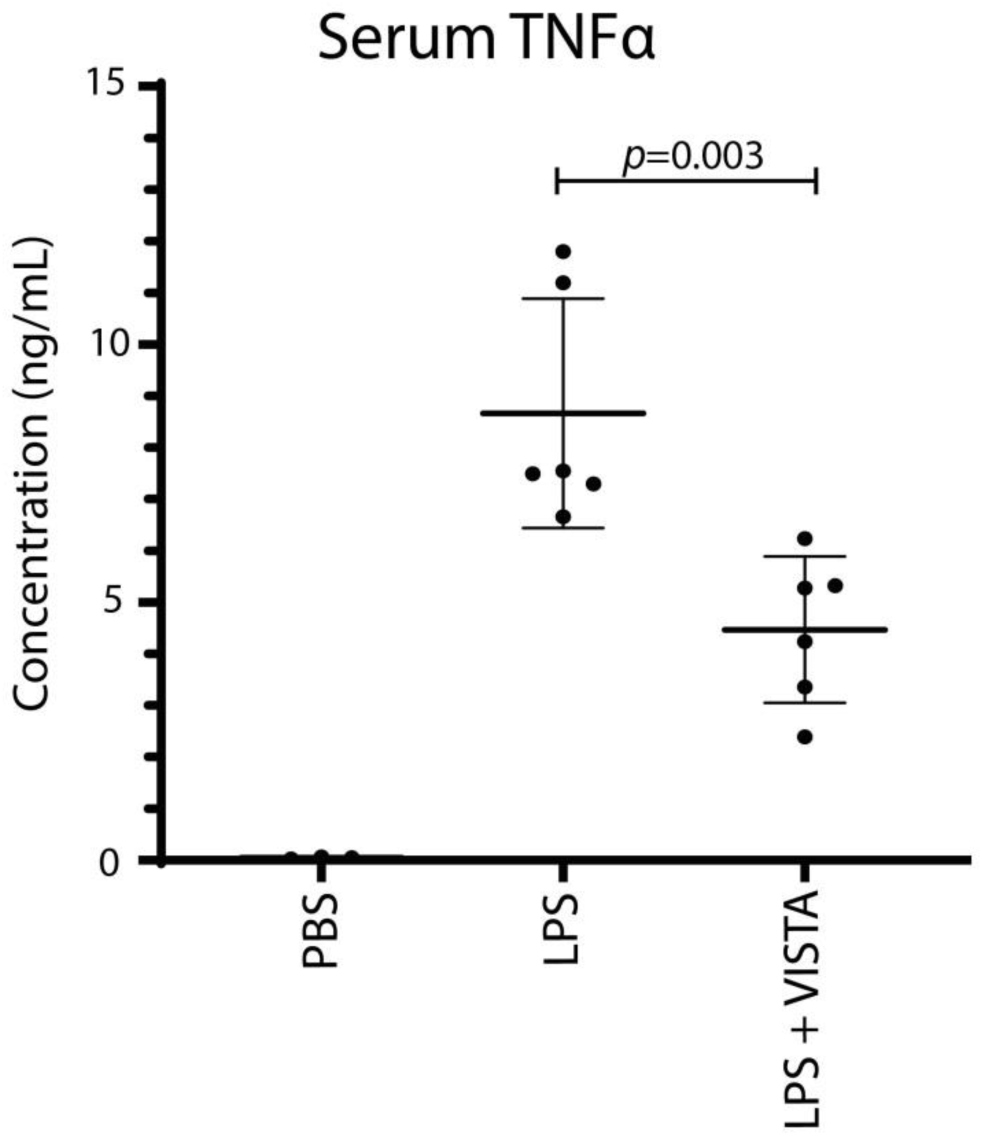
VISTA suppressed LPS-stimulated TNFα production *in vivo*. TNFα concentration in serum was analyzed 90 minutes after C57BL/6 mice were injected intraperitoneally with PBS, 10 mg/kg LPS, or 10 mg/kg LPS and 30 mg/kg VISTA-COMP.

## Discussion

VISTA has been proposed to function both as a ligand and as a receptor (1,2,4–8). In particular, agonistic anti-VISTA antibody was recently shown to reprogram macrophages to regulate innate immunity (7). Yet, the effect of VISTA behaving as a ligand on myeloid cells remains elusive. In this study, we demonstrated that exogenous VISTA acts as a ligand on macrophages and neutrophils to promote an anti-inflammatory profile by downregulating inflammatory cytokines and chemokines after LPS stimulation.

Interestingly, our findings parallel the dampening of pro-inflammatory cytokine/chemokine patterns observed when LPS-stimulated macrophages were treated with an agonistic anti-VISTA antibody (7). Specifically, both the presence of exogenous VISTA on LPS-stimulated peritoneal macrophages (Figures 2-4) and the use of anti-VISTA antibody on LPS-stimulated bone marrow-derived macrophages (7) reduce the expression of hallmark genes associated with LPS stimulation, *tnf* and *il6*. We also noted a downregulation in CXC chemokine gene signatures and protein secretion that are strongly associated with neutrophil chemotactic properties (33). Both treatments lead to the upregulation of immunoregulatory genes in LPS-stimulated macrophages, such as *lif* (28). These observations, combined with previous studies showing the similar immunosuppressive effect of both exogenous VISTA (1–5) and agonistic anti-VISTA antibodies (8,34,35) on T cells, suggests that VISTA acting as a ligand or a receptor evokes two redundant immunosuppressive pathways; potentially representing a self-regulation mechanism to control inflammatory responses. More studies in the context of VISTA as a self-regulation mechanism is warranted.

Interestingly, similar results were obtained when neutrophils were treated with VISTA-COMP. Neutrophils represent the most abundant circulating immune cell and are typically the first responders to several types of infection. If left uncontrolled, activated neutrophils will contribute to tissue damage during inflammatory processes poising them as targets for therapeutic intervention (36). The ability of exogenously added VISTA-COMP to dampen the inflammatory and chemotactic responses of neutrophils would thus suggest its use to treat acute inflammatory disorders.

As such, the therapeutic potential of VISTA-COMP in targeting activated myeloid cells in an acute inflammatory setting was tested in an *in vivo* model of endotoxemia. Macrophages and neutrophils are documented to play key roles in the imitation and progression of endotoxemia (30). In **Figure 6**, we showed that exogenously given VISTA-COMP significantly attenuated the increase in the characteristic monophasic spike in serum level of TNFα occurring upon LPS injection (26). This reduction is known to correlate with increased survival rates (32,37). These results further highlight VISTA-COMP as a potential immunotherapeutic agent to manage inflammatory events *in vivo*.

The nature of the VISTA receptor on macrophages and neutrophils remains an important question. Several candidates have been proposed by previous studies. VSIG3 has been identified as a ligand (38). However, it has not been shown to be expressed by hematopoietic cells, nor did our gene expression studies showVSIG3 to be expressed by resting or activated murine macrophages and neutrophils. Additionally, VISTA has been reported to bind to PSGL-1 in a pH-dependent manner (39). However, we have observed VISTA functioning at neutral pH, as similarly reported by another group (40), suggesting that the interacting partner of VISTA should be present and bind VISTA at neutral pH. Furthermore, like VSIG3, we did not find an increase in *psgl1* on activated murine macrophages and neutrophils. One key observation during this study, as shown in Figures 1 to 5, is that VISTA alone does not have an effect on un-stimulated cells. In contrast, significantly different gene signatures, implicated in downregulating inflammation, were observed when VISTA-COMP was added to LPS-stimulated cells. Our results suggest that the VISTA receptor is not constitutively expressed on resting macrophages and neutrophils, but is upregulated upon antigen (LPS in this case) stimulation. It would be interesting to determine whether other components linked to pathogen-associated molecular patterns (PAMPs) or damage-associated molecular patterns (DAMPs) would trigger similar upregulation of the receptor on myeloid cells. This knowledge may assist in further defining the receptor recognized by VISTA.

In conclusion, we have demonstrated a redundant cellular pathway where VISTA signals a receptor on myeloid cells to dampen the inflammatory immune response. In vivo results from an acute inflammatory model underscore the potential use of exogenous VISTA constructs such as VISTA-COMP as anti-inflammatory therapeutics.

