## Supplemental Files for "VISTA as a ligand downregulates LPS-mediated inflammation in macrophages and neutrophils"

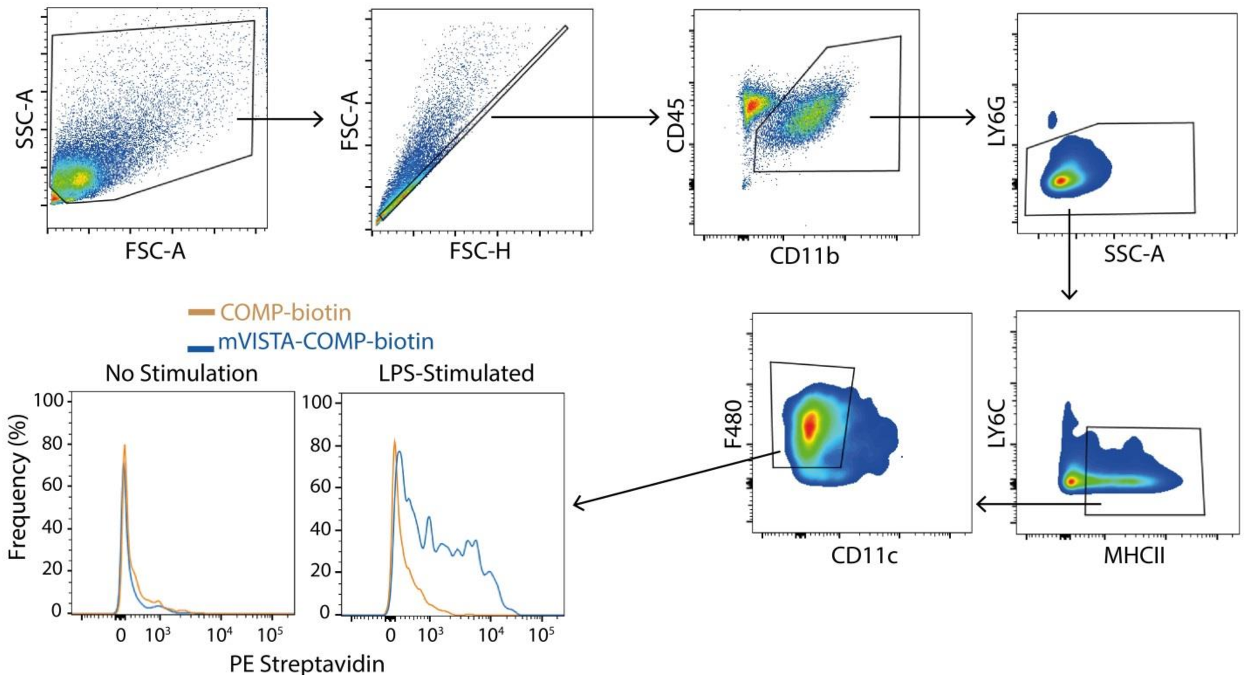


**Supplementary Figure 1: mVISTA-COMP binding to LPS-stimulated macrophages.** Flow cytometry was performed on PECs isolated 4 days after thioglycolate injection and stimulated with or without 1µg/mL LPS. PE-Streptavidin was used to detect the binding of biotinylated mVISTA-COMP or isotype control (biotinylated COMP) on macrophages.

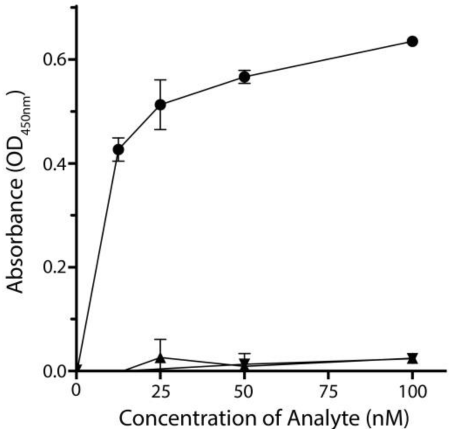


**Supplementary Figure 2: VISTA-COMP does not bind VISTA in a homotypic fashion.** To determine if VISTA binds homotypically, an ELISA was performed. Plates were coated with either VISTA-Fc or PD1-Fc (1 µg/ml) and VISTA-COMP or PDL1-COMP (5 µg/ml) binding was detected using an anti-his HRP-conjugated antibody and the substrate TMB. No binding was observed between VISTA-COMP and VISTA-Fc (▼) or VISTA-COMP and PD1-Fc (▲). As a positive control, PDL1-COMP was able to bind PD1-Fc (●). Absorbance readings were recorded at 450 nm. (n=3).

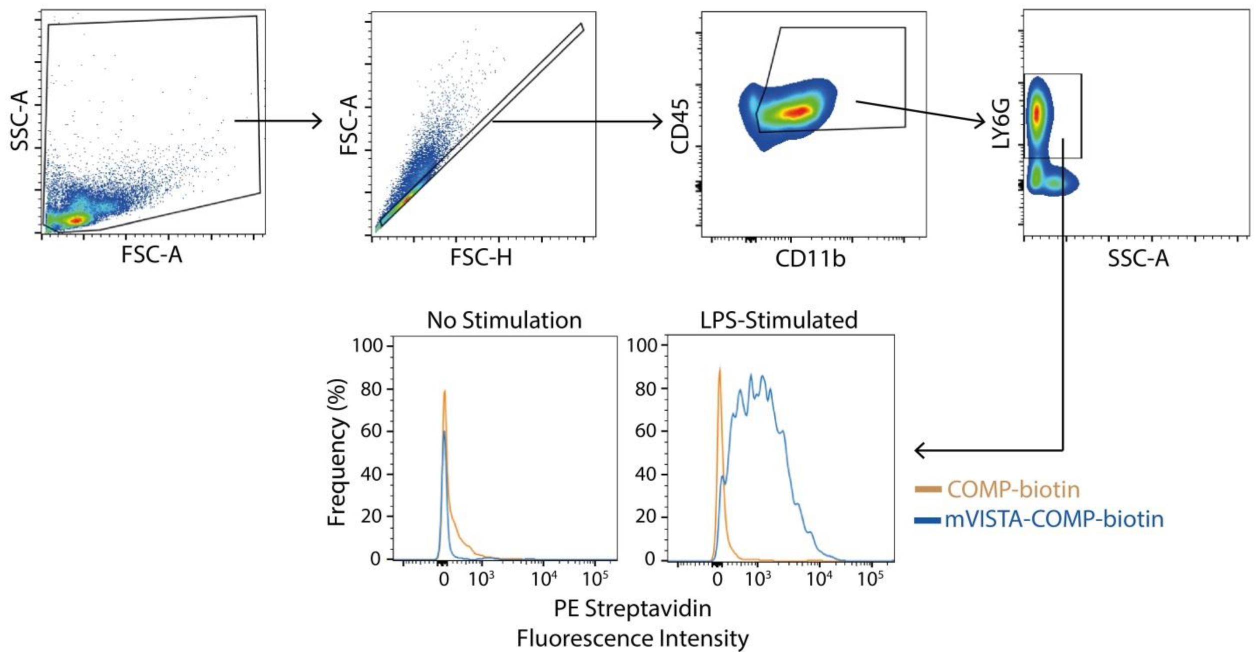


**Supplementary Figure 3: mVISTA-COMP binding to LPS-stimulated neutrophils.** Flow cytometry was performed on PECs isolated 1 day after thioglycolate injection and stimulated with or without 1µg/mL LPS. PE-Streptavidin was used to detect the binding of biotinylated mVISTA-COMP or isotype control (biotinylated COMP) on neutrophils.
